# Combining functional and structural genomics to track antibiotic resistance genes in mobile elements in clinical bacterial strains

**DOI:** 10.1101/2020.10.30.361923

**Authors:** Tiago Cabral Borelli, Gabriel Lencioni Lovate, Ana Flavia Tonelli Scaranello, Lucas Ferreira Ribeiro, Livia Zaramela, Felipe Marcelo Pereira-dos-Santos, María-Eugenia Guazzaroni, Rafael Silva-Rocha

## Abstract

The rise of multi-antibiotics resistant bacteria represents an emergent threat to human health. Here, we investigate antibiotic resistance mechanisms in bacteria of several species isolated from an intensive care unit in Brazil. We used whole-genome analysis to identify antibiotic resistance genes (ARGs) and plasmids in 35 strains of Gram-negative and Gram-positive bacteria, including the first genomic description of *Morganella morganii* and *Ralstonia mannitolilytica* clinical isolates from South America. We identify a high abundance of beta-lactamase genes in highly resistant organisms, including seven extended-spectrum β-lactamases shared between organisms from different species. Additionally, we identify several ARGs-carrying plasmids indicating the potential for fast transmission of resistance mechanism between bacterial strains, comprising a novel IncFII plasmid recently introduced in Brazil from Asia. Through comparative genomic analysis, we demonstrate that some pathogens identified here are very distantly related to other bacteria isolated worldwide, demonstrating the potential existence of endemic bacterial pathogens in Brazil. Also, we uncovered at least two couples of (near)-identical plasmids exhibiting multi-drug resistance, suggesting that plasmids were transmitted between bacteria of the same or different species in the hospital studied. Finally, since many highly resistant strains carry several different ARGs, we used functional genomics to investigate which of them were indeed functional. In this sense, for three bacterial strains (*Escherichia coli*, *Klebsiella pneumoniae*, and *M. morganii*), we identify six beta-lactamase genes out of 15 predicted *in silico* as the main responsible for the resistance mechanisms observed, corroborating the existence of redundant resistance mechanisms in these organisms.

**Importance:** Big data and large-scale sequencing projects have revolutionized the field, achieving a greater understanding of ARGs identification and spreading at global level. However, given that microbiota and associated ARGs may fluctuate across geographic zones, hospital-associated infections within clinical units still remain underexplored in Brazil – the largest country in South America; 210 million inhabitants – and neighboring countries. This work highlighted the identification of several ARGs shared between species co-occurring simultaneously into a Brazilian hospital, some of them associated with large plasmids, mostly endowed with transposable elements. Also, genomic features of clinically underrepresented pathogens such *M. morganii* and *B. cepacia* were revealed. Taken together, our results demonstrate how structural and functional genomics can help to identify emerging mechanisms of shared antibiotic resistance in bacteria from clinical environments. Systematic studies as the one presented here should help to prevent outbreaks of novel multidrug resistance bacteria in healthcare facilities.

## Introduction

Microbial resistance to antibiotics is a growing global concern. Established protocols in clinics to fight nosocomial infections include isolation of microorganisms from patient samples to allow its identification and to determine its antibiotic susceptibility (1). However, this process is time-consuming (48 h or more) and prone to pathogen misidentification (2). Therefore, the advent of next-generation sequencing (NGS) tools has allowed the rise of novel approaches to identify microbial pathogens and to fight infection (3). Thus, in the last two decades, whole-genome sequencing (WGS) of microbial pathogens has moved from being used as a basic research tool to understand pathogen’s biology and evolution (4, 5), to an almost routine diagnostic tool to investigate outbreaks in hospitals and nosocomial infection pathways (6–9). For diagnostics purposes, current WGS technologies could even be cost effective for slow-growing pathogens such as *Mycobacterium tuberculosis*, providing faster and accurate results even for antibiotic resistance determination (10). Additionally, culture independent methods based on clinical metagenomics can be used to identify several pathogens from nucleic acids extracted from patient samples without the need for microbial isolation (11–13). Furthermore, recent progress on the use of artificial intelligence tools has allowed the construction of computational models that can predict with high accuracy antimicrobial susceptibility of microbial pathogens based on WGS data (14–16).

While WGS analysis as routine for microbial identification is not a worldwide reality, it has been extensively used to investigate microorganisms’ population structure at different scales. For example, Arias and coworkers using WGS analysis from 96 methicillin-resistant *Staphylococcus aureus* (MRSA) from 9 countries in Latin America demonstrated a high degree of variation in the genome of different isolates from those countries (17). The same study indicated that among sampled hospitals, those in Brazil presented a higher incidence of MRSA strains (up to 62%). Similarly, a recent work by David and coworkers investigated the path for the nosocomial spread of *Klebsiella pneumoniae* in 244 hospitals in 32 European countries (18). Using a well-defined sampling strategy and WGS analysis of more than 1700 *K. pneumoniae* strains, authors were able to quantify the role of intra-hospital pathogen dissemination, as well as some potential paths for the introduction of novel strains from the USA to Europe. In addition to those examples, large-scale WGS analysis has been used to investigate the molecular adaptation to different hosts, as in the work by Arimizu *et al.* in which authors analyzed *Escherichia coli* strains from human versus bovine samples (19).

Another key process playing a significant role in the rise of new microbial threats is the propagation of mobile virulence and antimicrobial resistance factors within these populations mediated especially by plasmids and transposons (5, 20). Therefore, the rapid evolution of plasmids through structural rearrangements, virulence genes acquisition, plasmid fusions, and propagation to pathogens can account for the fast dissemination of supervirulent or super-resistant bacteria (21–23). Understanding the very dynamic and complex processes could hold the potential to design new drugs aiming at reducing plasmid propagation between pathogens (24). While the use of WGS is currently growing worldwide, most studies (especially in Brazil) have been restricted to some particular species (such as *K. pneumoniae* or *S. aureus*) without considering their interplay with other species in hospital settings. Here, we investigate at the genomic level 35 strains from 18 different species (and 11 genera) isolated at the same two weeks in a reference public hospital in Brazil. WGS analysis indicates that many strains are only distantly related to those available at public databases. We aimed to identify antibiotic resistance genes (ARGs) and plasmids harboring these elements, as well as evidence for common resistance mechanisms shared between strains from the same or different species. We were able to identify several ARG-harboring plasmids, two of them present in both Gram-negative and Gram-positive strains, and seven beta-lactamases located in multiple hosts with 100% identity at the nucleotide level, two of which were inferred to be active using functional genomic library screening. In addition, comparative sequence analysis identified a novel IncFII *K. pneumoniae* plasmid harboring two ARGs, potentially indicating a recent introduction from Asia to Brazil.

## Results and discussion

### WGS analysis of clinical strains isolated from the same two weeks

We selected 35 bacterial strains isolated from different patient samples and performed WGS as represented in **Fig. 1**. While well-studied pathogens such as *K. pneumoniae*, *E. coli*, *P. aeruginosa* e *S. aureus* were well represented in the sampling, we were able to analyze pathogens with very few representative genomic information in March 2020. For example, two *Ralstonia mannitolilytica* strains were sequenced, but only nine complete or graft genomes were available at NCBI. Other underrepresented strains were *Streptococcus gallolyticus* (27 genomes available) and *Morganella morganii* (63 genomes). These numbers contrast with those from *K. pneumoniae*, *E. coli,* or *S. aureus,* where 8-20 thousand genome sequences are available. This evidence indicates that many clinically relevant pathogens have been underrepresented in WGS analysis efforts worldwide. For instance, phylogenetic analysis of *Burkholderia cepacia* 540A against all 166 available genomes in NCBI (**Fig. S2**) indicates this strain is also very divergent but belongs to a branch formed by strains isolated from patients with cystic fibrosis in the UK and endophytic bacteria isolated from Australia (25). Therefore, some of the new WGSs generated here demonstrate a significant diversity of some underrepresented microorganisms and should serve as reference sequences for future studies on clinical isolates in Brazil and South America.

**Figure 1.**
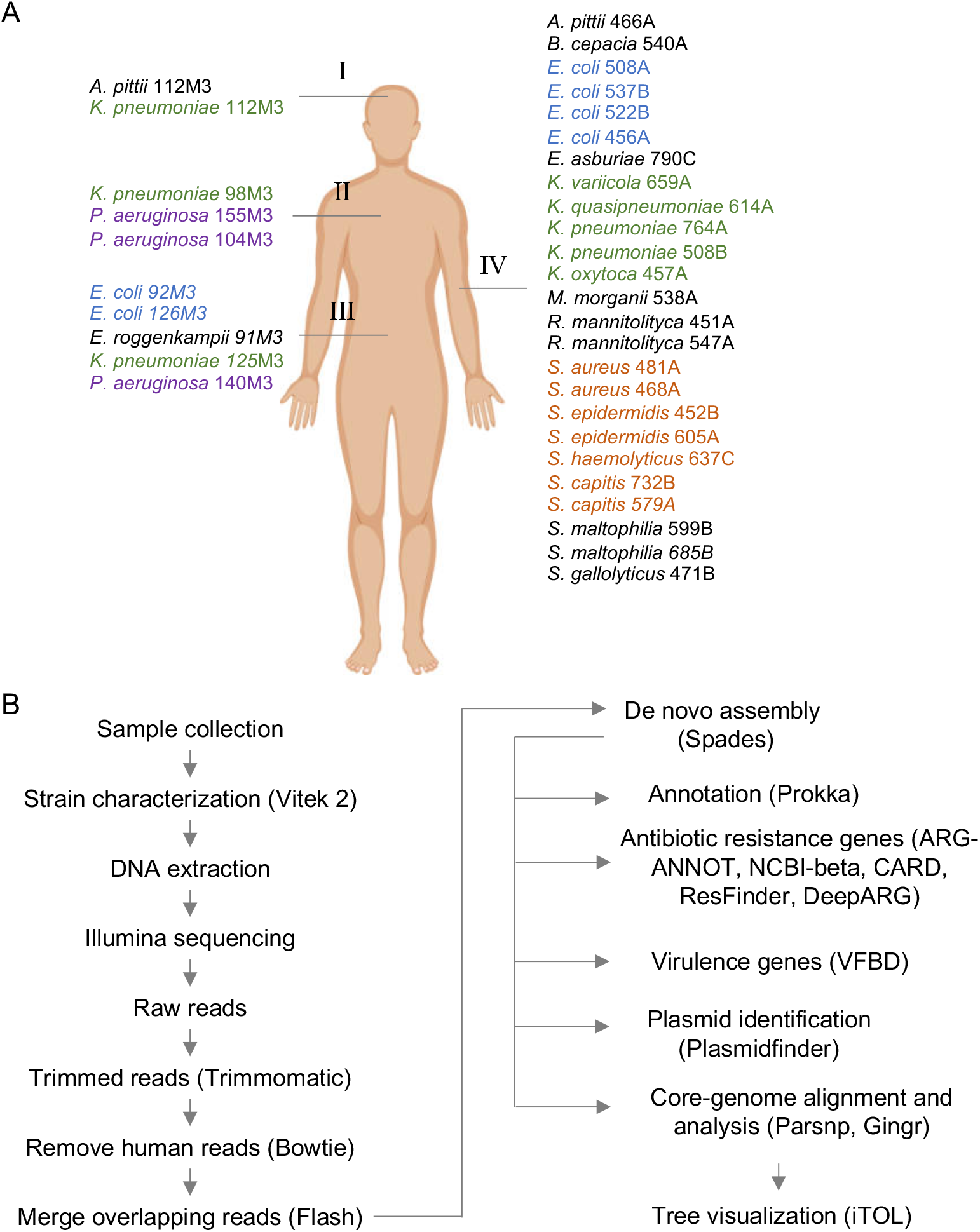
Overall strategy used for the whole-genome analysis of clinical strains. **A**) In total, 35 bacterial strains were isolated from several samples, such as cerebrospinal fluid (I, 2 strains), bronchoalveolar lavage (II, 3 strains), ascitic fluid (III, 5 strains), and blood (IV, 25 strains). The four most common bacterial groups (*K. pneumoniae*, *E. coli*, *P. aeruginosa,* and *S.* genus) are colored. **B**) Schematic representation of the main bioinformatic pipeline used for genome sequencing, assembly, annotation, identification of ARGs, and virulence factors and genomics analysis.

### Identification of resistance genes in clinical strains

We next analyzed the existence of ARGs in the sequenced genomes. Analysis of ARGs using ARG-ANNOT database indicated a higher prevalence of beta-lactamase coding genes followed by amino-glycosidases (**Fig. 2A**), while analysis with DeepARG tool (26) showed the multidrug category as the most abundant on the genomes analyzed (**Fig. S3**). We next analyzed ARGs’ distribution in the two most abundant groups, *E. coli* and *Klebsiella*. In the first case, each of the six *E. coli* strains displays a unique set of ARGs from different categories, and five particular beta-lactamases (*bla*_TEM-105_, *bla*_OXA-1_, *bla*_KPC-2_, *bla*_CTX-M-15,_ and *bla*_CMY-111_) were found in at least one genome (**Fig. 2B**). Only one strain (*E. coli* 456A) did not present any of these five *bla* genes, and this strain was sensitive to all antibiotics tested by Vitek 2 (**Fig. 2B**/**Fig.3**). For the *Klebsiella* group, we observed a much broader set of resistance markers (**Fig. 2C**), which was in accordance with the largest level of antibiotic resistance of this group compared to *E. coli*. Next, we investigated which beta-lactamase genes were conserved into the strains analyzed. For this, we compared all ~125 *bla* genes identified in **Fig. 2** and searched for those coding proteins with 100% identity at the amino acid sequence (**Fig. 3**). Using this approach, we identified seven *bla* genes (*bla*_OXA-1_, *bla*_OXA-10_, *bla*_CTX-M-1_, *bla*_KPC_, *bla*_TEM_, *bla*_HYDRO,_ and *bla*_BLP_) which have 100% aa identity between two or more strains. Notably, *bla*_OXA-1_, *bla*_CTX-M-1_, *bla*_KPC,_ and *bla*_TEM_ were present in 3 to 6 strains from different species. Interestingly, both *bla*_OXA-1,_ and *bla*_CTX-M-1_ were found in *E. coli*, *K. pneumoniae* and *M. morganii*, which could reveal recent horizontal gene transfer between these strains. From these seven genes, 2 (*bla*_OXA-1_ and *bla*_KPC_) were functional in the library screening presented below. Taken together, since the analyzed strains were isolated from hospitalized patients in the same two weeks, this evidence would indicate a recent mobilization of antibiotic resistance determinants either in the environment or in the hospital settings, as was reported previously (27–29).

**Figure 2.**
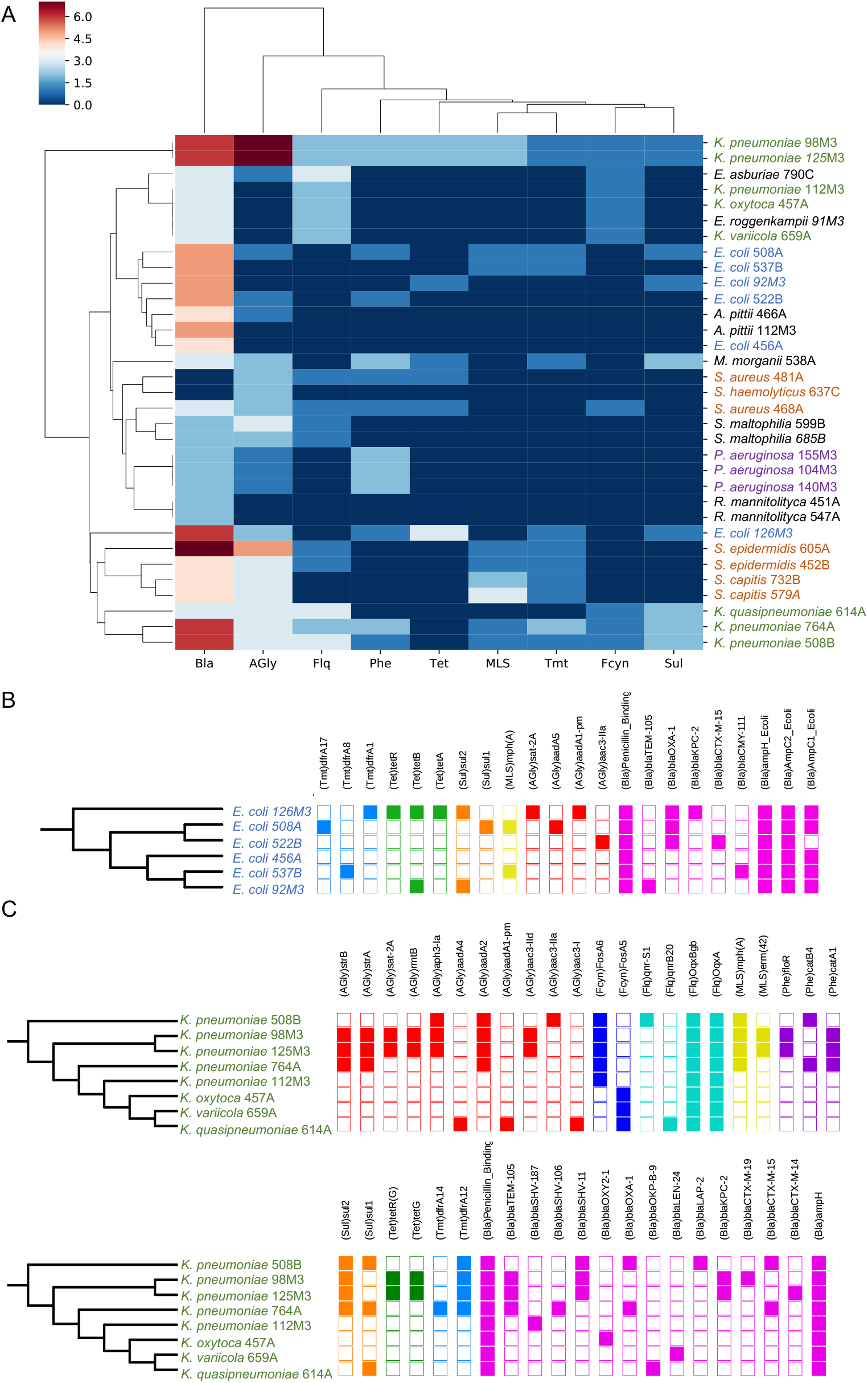
Identification of ARGs in nearly sequenced genomes. **A**) Heatmap showing the presence of ARGs identified by ARG-ANNOT. Only *B. cepacia* did not present any identified ARG. Data was clustered using hierarchical mapping with Euclidian distance. Blue to red scale indicate number of ARG for each strain in each category, as indicated in the legend. **B**) Distribution of different ARGs per genome of *E. coli*, colored by antibiotic category. The maximum likelihood phylogeny for the strains was based on core-genome. **C**) Distribution of different ARGs per genome of *K. pneumoniae*, following the scheme in B.

**Figure 3.**
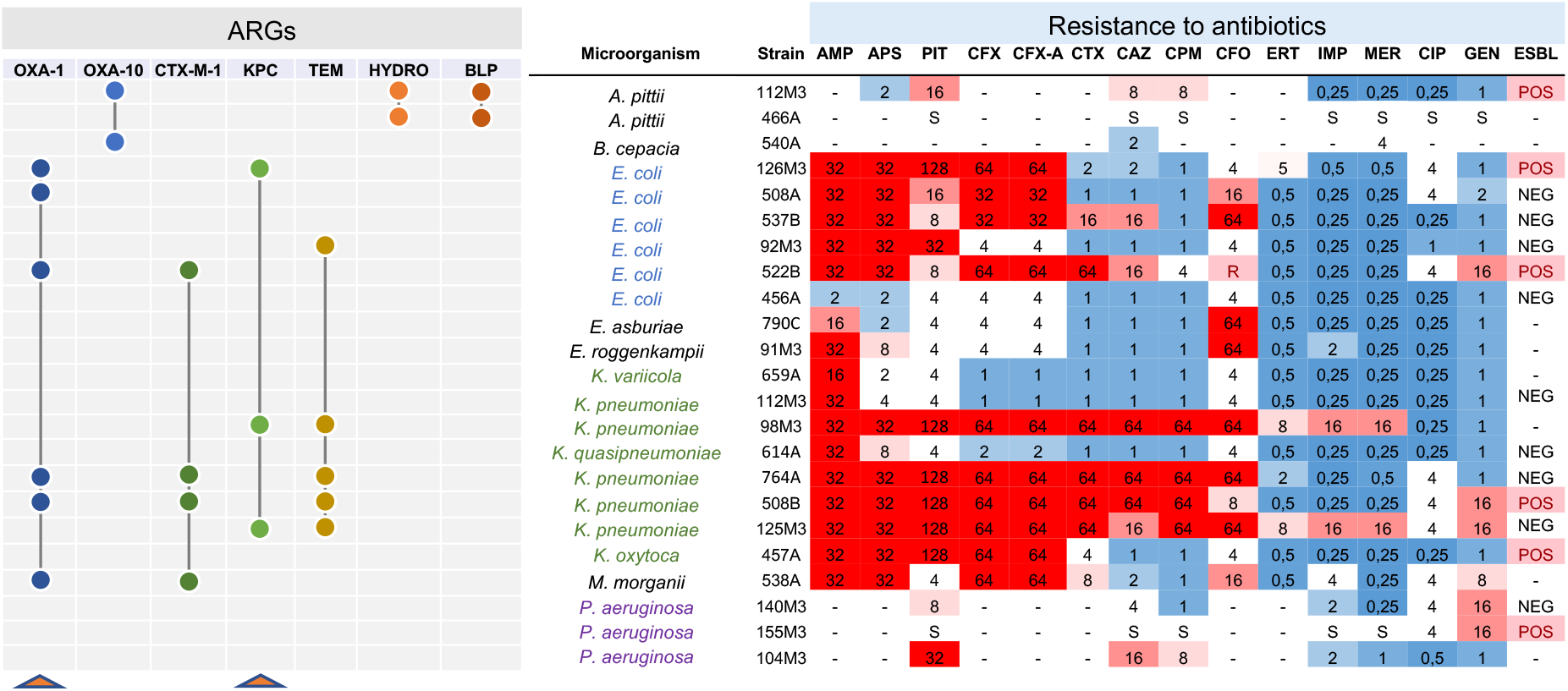
Shared beta-lactamases with 100% identity at the protein level. Seven beta-lactamase coding genes were found as shared among gram-negative strains analyzed. Connected circles indicate that the genes are presented in those strains. On the right, the antibiotic resistance profile of the analyzed strains. Genes for *bla*_OXA-1_ and *bla*_KPC_ are highlighted (red triangles) since these genes were identified in the functional screening carried out in this study. On the right, antibiotic resistance levels are indicated, with numbers indicating the resistance levels in μg/mL. Red indicates resistance to the antibiotic while blue denotes sensibility. The final column indicated if the strain is ESBL positive or negative.

### Identification of ARGs located in plasmids

We next aimed to identify ARGs with potential mobilization through plasmids in the analyzed species. For this, we crossed the data from ARG-ANNOT with the prediction of plasmid elements generated by PlasmidFinder. Using this approach, we were able to identify nine potential plasmids from 7 species associated with at least one ARG. As shown in **Fig. 4A**, a ~42kb plasmid –(pKP98M3N42) – harboring a *bla*_KPC-2_ and a *sat-2A* resistance determinant were identified in *K. pneumoniae* 98M3, and this plasmid also carries transposases, recombinases, and type IV secretion system genes. An identical plasmid – (pKP125M3N44; 100% nucleotide sequence identity) – was also found in *K. pneumoniae* 125M3 (**Fig. 4A, Fig. S4A**). As mentioned before, this *bla*_KPC-2_ gene was identical in these two *K. pneumoniae* strains and in *E. coli* 126M3, but it was not located in a plasmid in the latter case (**Fig. 3**). In two recent studies, authors demonstrate ARG-carrying plasmids transfer in clinical settings between bacteria of the same or different species, which emphasize the significance of revealing potential plasmid-mediated outbreaks to efficiently track ARGs horizontal transmission in hospitals (30, 31).

**Figure 4.**
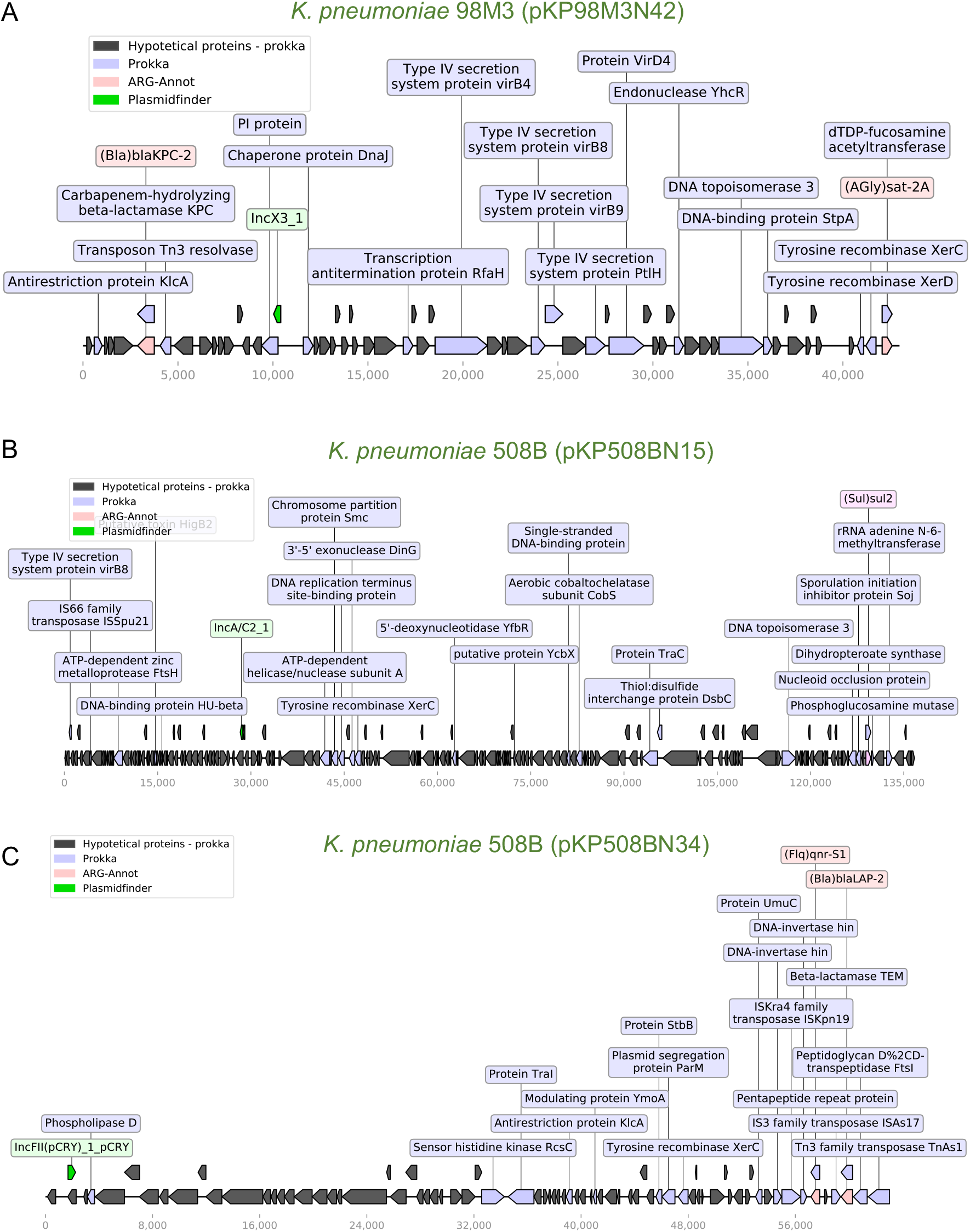
Schematic representation of genes of three *K. pneumoniae* plasmids. **A**) Plasmid pKP98M3N42 (43 kb) from *K. pneumoniae* 98M3. This plasmid carries two ARGs (*bla*_KPC-2_ and *sat-2A*), elements of a type IV secretion system, two resolvases, and a Tn3 transposase. This plasmid is very similar to pKP125M3N44 from *K. pneumoniae* 125M3 (Fig. S4A). Whole-plasmid visualization was performed using a python module for prokaryotic genome analyses – DnaFeaturesView – and matplotlib module combined, as well as ARG-ANNOT and Prokka’s results (57).**B**) Plasmid pKP508BN15 (136.8 kb) from *K. pneumoniae* 508B transports a *sul2* and an rRNA adenine n-6-methyltransferase (*ermC*) resistance markers. A transposase, a *xerC* recombinase, and a type IV secretion system *virB8* protein are also found in this plasmid. **C**) Plasmid pKP508BN34 (63 kb) is also from *K. pneumoniae* 508B and carries several transposases and two resistance determinants, *qnr-S1* and *bla*_LAP-2_. Legends represent the colors code for the identified genes.

Another strain, *K. pneumoniae* 508B, harbors two plasmids with ~136kb (pKP508BN15) and ~62kb (pKP508BN34). Both plasmids harbor transposon elements, with the larger one harboring a *sul2* resistance gene and the smaller one with two ARGs (*qnr-S1* and *bla*_LAP-2_, **Fig. 4B-C**). In general, ARGs’ coexistence with transposon elements was also observed for two plasmids identified in *E. coli* strains (**Fig. S4B-C**) and *Staphylococcus capitis* 732B (**Fig. S5A**). Finally, two almost identical small plasmids (~2.3kb; 99.94% nucleotide identity) were identified in *S. capitis* 732B and *S. epidermidis* 452B, which harbor an *aadC* and *ermC* resistance determinants (**Fig. S5B**). Taken together, these data demonstrate the potential for dissemination of many ARGs genes identified here, and the existence of identical or near-identical plasmids between different species could indicate that these elements have been mobilizing among some of these species.

### Experimental validation of functional beta-lactamases from highly-resistance bacteria

Once we distinguish several ARGs in the genomes analyzed, we decided to perform a functional screening to identify which of these genes could confer resistance into a heterologous host. For this, we selected three strains (*E. coli* 126M3, *K. pneumoniae* 508B, and *M. morganii* 538A) to construct genomic libraries into laboratory *E. coli* DH10B (Table 1). The libraries were constructed into the broad host range vector pSEVA232, which contains a kanamycin resistance marker, a broad host range *oriV* with a medium-copy number, and a variant of the *lacZα* multiple cloning site with a *Plac* promoter (32–34). In this way, the libraries generated during this study or individual plasmids of interest can be transferred to other bacterial strains for another screening or functional evaluation (**Fig. 5A**). The use of a medium copy-number plasmid allows a closer assessment of ARGs’ natural genetic context in contrast to other studies that use high copy plasmids (35). Accordingly, plasmids with lower copy-number and monomeric states also tend to be more stably inherited throughout bacterial populations (36). We screened ~750.000 clones of each library against each antibiotic (amoxicillin, oxacillin, and penicillin G) and obtained 44 clones with unique sequences containing ARGs (**Fig. 5B** and **Table 2**).

**Table 1.**
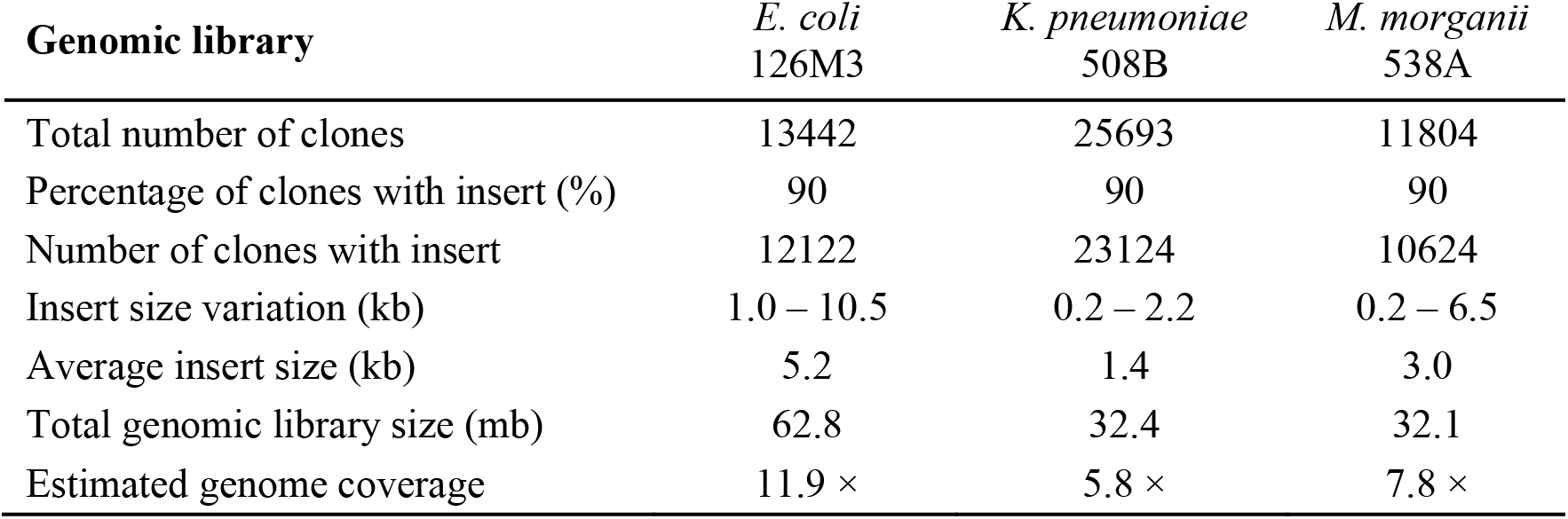
Features of the generated genomic libraries.

**Table 2.** Comparison of ARGs identification using *in silico* and functional approaches.

| Strain | *In silico* (argannot) | No. of sequenced resistant clones |
| --- | --- | --- |
| *E. coli* 126M3 | Penicillin_Binding_Protein_Ecoli | - |
|  | AmpC1_Ecoli | - |
|  | *bla*<sub>KPC-2</sub> | 28 |
|  | AmpC2_Ecoli | 1 |
|  | *bla*<sub>OXA-1</sub> | - |
|  | ampH_Ecoli | - |
| *K. pneumoniae* 508B | *bla*<sub>SHV-11</sub> | - |
|  | Penicillin_Binding_Protein_Ecoli | - |
|  | *bla*<sub>LAP-2</sub> | 2 |
|  | ampH | - |
|  | *bla*<sub>CTX-M-15</sub> | 8 |
|  | *bla*<sub>OXA-1</sub> | 1 |
| *M. morganii* 538A | *bla*<sub>CTX-M-15</sub> | - |
|  | *bla*<sub>OXA-1</sub> | - |
|  | *bla*<sub>MOR-2</sub> | 4 |

**Figure 5.**
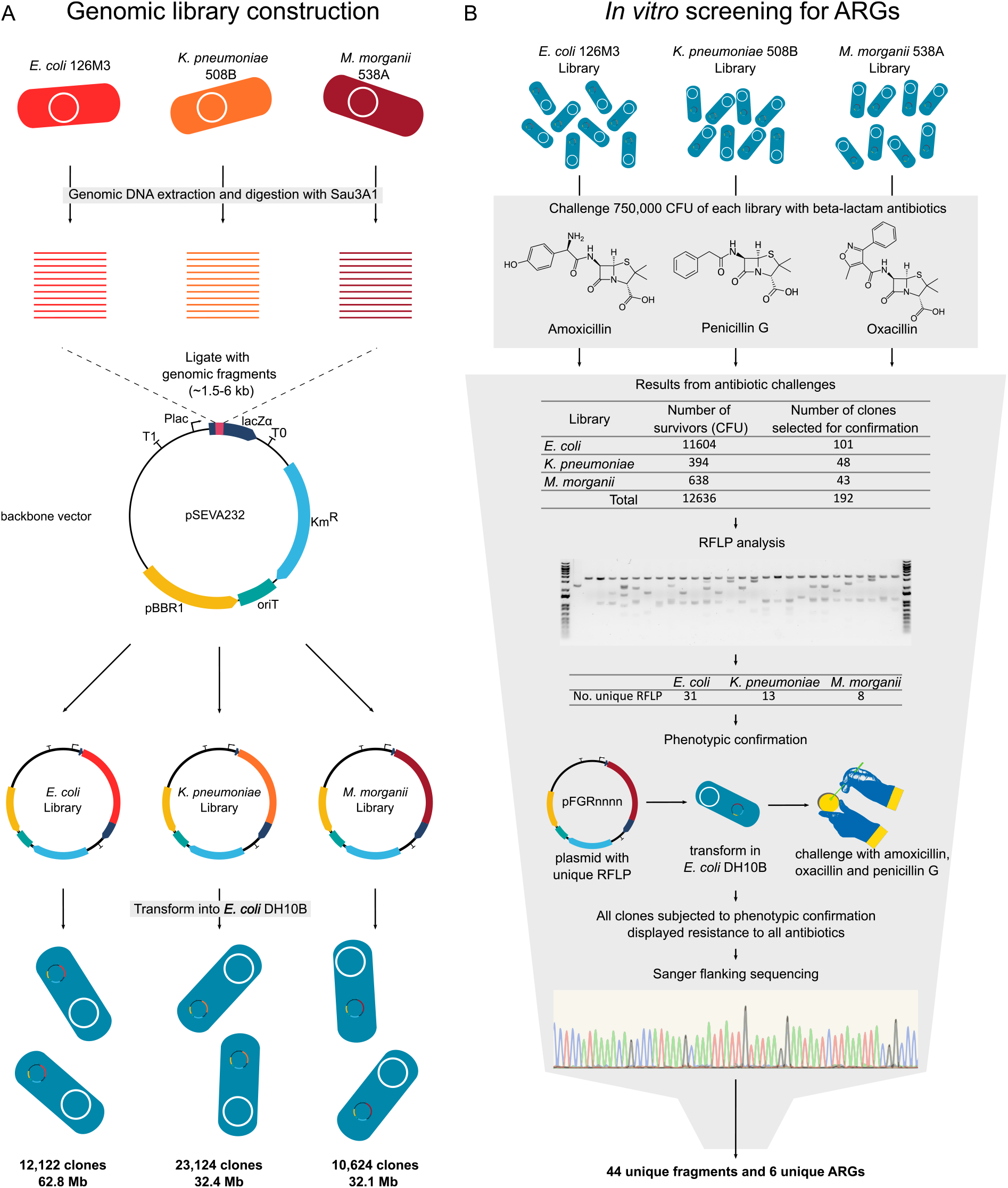
Experimental design for functional genomics analysis. **A**) Strains and backbone used to construct the library and features of the obtained libraries. **B**) Functional screening for beta-lactam resistant clones, phenotypic confirmation, and sequence identification. CFU – colony forming unit, RFLP - restriction fragment length polymorphism, KmR - kanamycin resistance marker, pBBR1 - a broad host range *oriV*, *lacZα* - *lacZα* gene with multiple cloning site, *Plac* - *Plac* promoter, pFGRnnnn – plasmid naming schema.

Clones containing the *bla*_KPC-2_ were by far the more abundant in the screening (**Table 2** and **Fig. 6**). The genomic context of identified *bla*_KPC-2_ indicate that it is prone to suffer horizontal transfer, once it is flanked by transposases. Martínez *et al.* (37) describe a framework to prioritize the risk of ARGs, the Resistance Readiness Condition (RESCon). The RESCon algorithm considers the similarity of an ARG to known genes, functional evaluation, the clinical relevance of the antibiotic, presence of a mobile genetic element, and presence in a human pathogen. The characteristics of this ARG would categorize it as RESCon 1, an ARG with the highest possibility to thrive in a clinical setting (37). Although a framework better describing the impacts of gene transfer to prioritize risk is needed (38), the RESCon classification indicates clinical relevance of this ARG. With ARGs found in all 3 functional screenings, we were able to identify with high frequency six different *bla* genes using the functional approach presented here, being at least two of them present in plasmids – *bla*_KPC-2_ and *bla*_LAP-2_; see next section – (**Fig. 6**).

**Figure 6.**
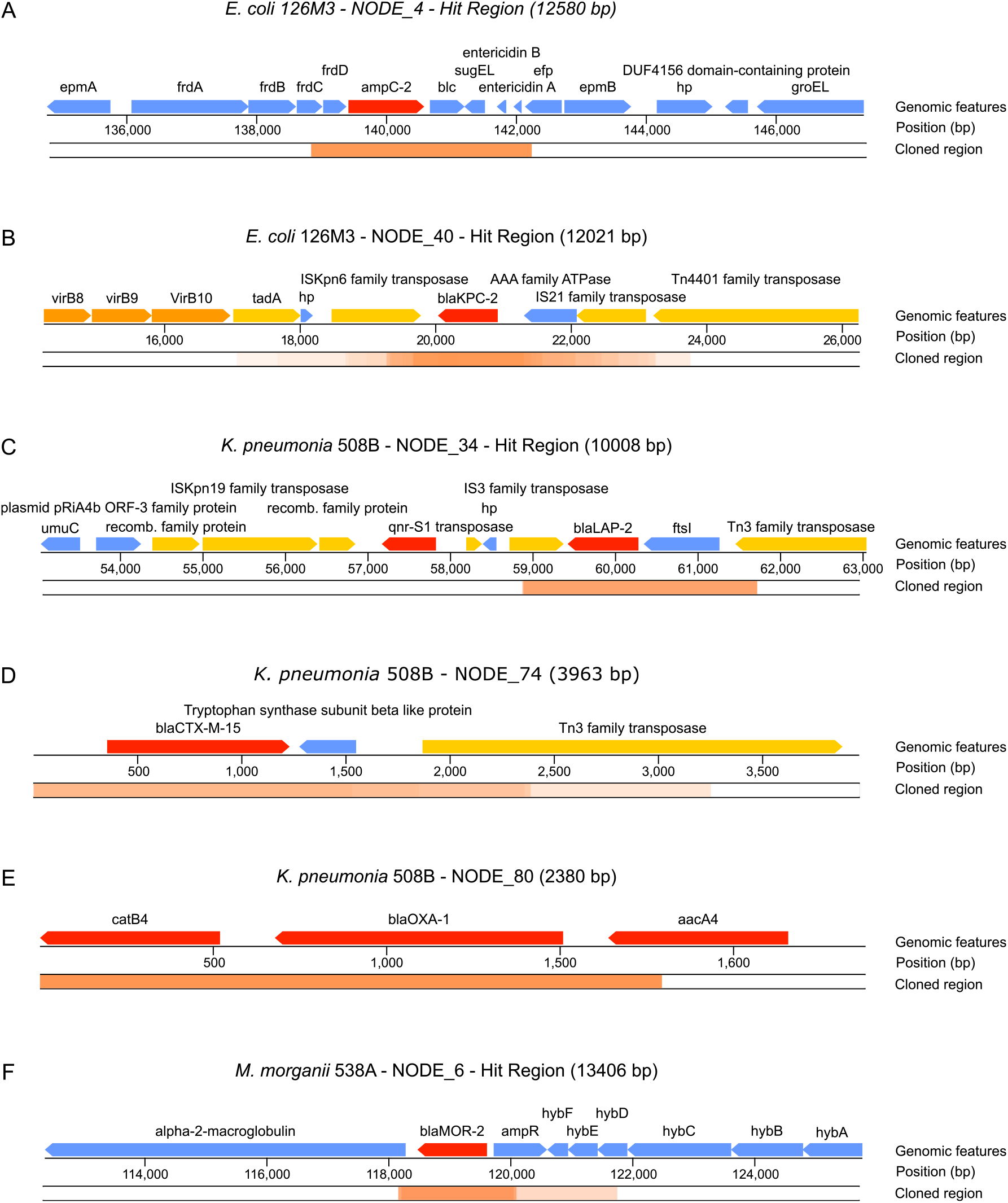
Features found in identified clones conferring resistance to antibiotics. **A**) *ampC-2* gene identified from *E. coli* 126M3. **B**) *bla*_KPC-2_ gene identified from *E. coli* 126M3. **C**) *bla*_LAP-2_ gene identified from pKP508BN34 plasmid from *K. pneumoniae* 508B. **D**) *blaCTX-M-15* gene identified from *K. pneumoniae* 508B. **E**) *bla*_OXA-1_ gene identified from *K. pneumoniae* 508B. **F**) *bla*_MOR-2_ gene identified from *M. morganii* 538A. ORFs were colored according to its function: red – ORFs directly related to antibiotic resistance; orange – ORFs related to virulence; yellow – ORFs related to horizontal transfer of the ARG; blue – ORFs with no identified relation to pathogenesis. The cloned region represents contig regions that were identified in our screenings. Overlapping cloned regions are darker, while dim regions have fewer overlapping identified clones.

We propose that functional genomics can be combined with current approaches based on large-scale sequencing in order to better understand the functional aspects of ARGs. Recent developments in machine learning (26) have provided tools to find ARGs that can be used to detect ARGs that would not be otherwise identified with sequence similarity tools. Still, the databases used to train those tools are biased for specific antibiotic resistance classes, such as beta-lactam, bacitracin, MLS, and efflux-pumps. Indeed, we could only find just a fraction of ARGs annotated through bioinformatics tools with our functional approach. This paradox could be due to the other ARGs not being functional in the experimental conditions used here or because our screening was not exhaustive enough to cover those sequences.

### Mapping of plasmids in the Brazilian territory

To evaluate the biogeographic distribution of functional ARGSs from *K. pneumoniae*, we performed genomic comparative analysis including 50 plasmids deposited in NCBI data bank and two of the plasmids identified in this study (pKP98M3N42, pKP508BN34). Distance-tree analysis showed the presence of two distinguishable groups. Plasmid pKP98M3N42 (identical to plasmid pKP125M3N44, also identified in this work) is located in the first group – comprising seven strains all reported in Brazilian cities from 2009 to 2015, but 1 in USA –, which should indicate that pKP98M3N42 is sharing structural features with the other plasmids positioned in this group (**Fig. 7A**). Additionally, pKP98M3N42 shares 100-99.9% sequence identity with IncX3 plasmids of these seven strains that have been previously related to play an important function in mediating horizontal transmission of *bla*_KPC-2_ genes among hospital-associated members of the Enterobacteriaceae family (39, 40). Moreover, IncX3 plasmids carrying *bla*_KPC-2_ genes were also reported in countries very distant geographically, such as United States (41), Brazil (42, 43), Australia (44), Italy (45), France (46), South Korea (47, 48), China (49), Israel and Greece (39), to cite some. Interestingly, although IncX3 self-transmissible plasmids are widespread globally, the evident diversification of plasmids in the first separated branch of the tree (**Fig. 7A**) could indicate that plasmids from strains isolated in clinically relevant bacteria in Brazilian ground undergo their own structural reorganization.

**Figure 7.**
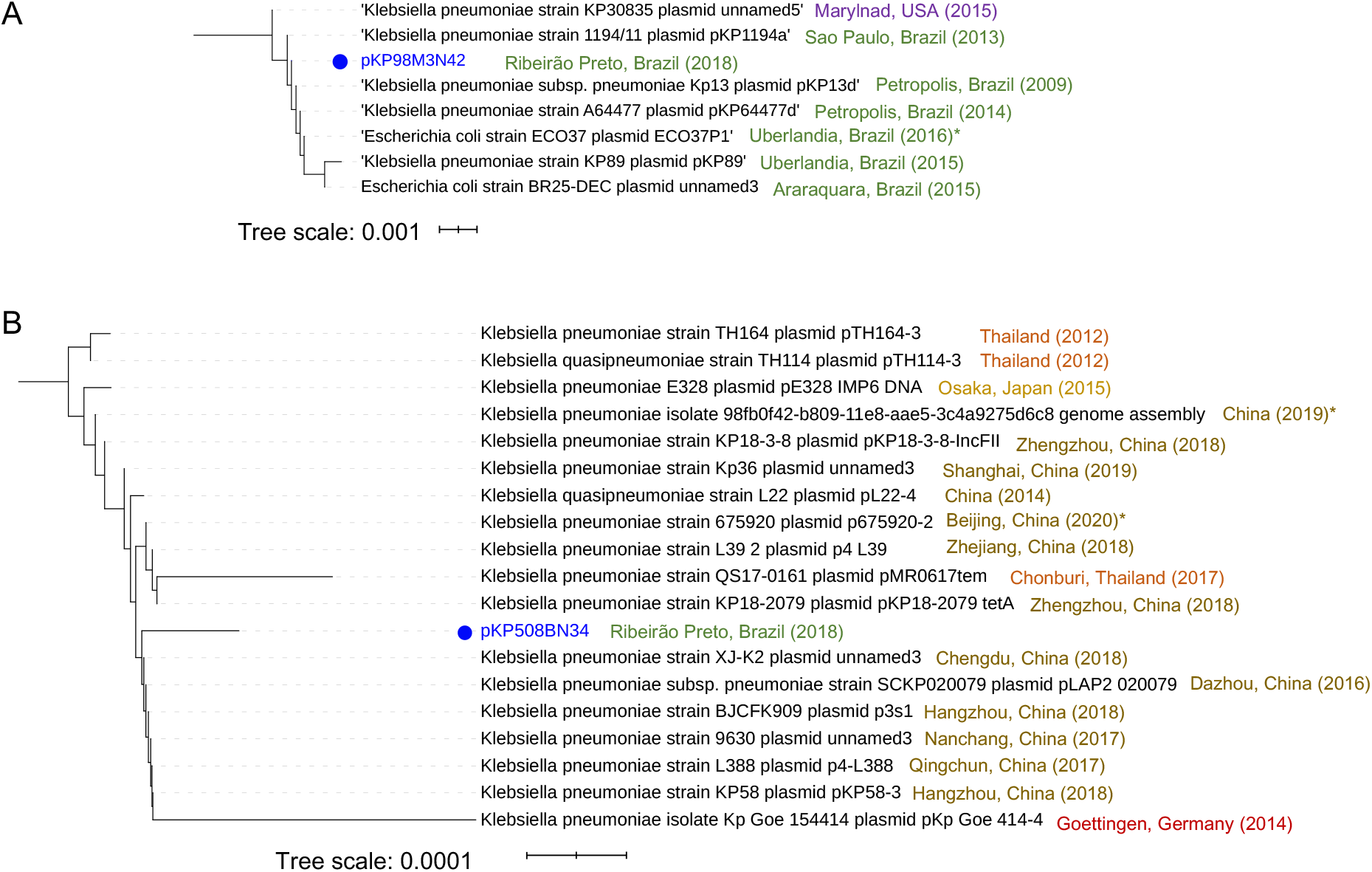
Distance relationship analysis for two plasmids from *K. pneumoniae*. **A**) Distance relationship of the 7 closest plasmids’ nucleotide sequences to pKP98M3N42 showing an *E*-value of less than 0.0 and a minimal sequence cover of 70% in BLAST analysis. The tree was produced using pairwise alignments by means of the Fast-minimum evolution method. Year denotes the collection data and the asterisk indicates the data reported in the public database. **B**) Distance relationship of the 18 closest plasmids’ nucleotide sequences to pKP508BN34 showing an *E*-value of less than 0.0 and a minimal sequence cover of 50% in BLAST analysis. The tree was produced using pairwise alignments by means of the Fast-minimum evolution method. Year states the collection data and the asterisk indicates the data reported in the public database, informed when collection data was not available. iTOL (https://itol.embl.de) was used for tree visualization.

On the other hand, distance tree representation of the 50 close plasmids’ sequences related to pKP508BN34 did not show the presence of evident different groups. BLAST analysis showed that plasmid pKP508BN34 is widely distributed in *K. pneumoniae* strains, with 100-99,9% sequence identity and with query coverage ranging from 84% to 100% to the 18 closest related plasmids, some of them associated to hypervirulent strains – such as strain KP58, bearing plasmid pKP58-3 – (**Fig. 7B**). Mortality due to infection of *K. pneumoniae* was very rare in the past. However, this pathogen’s fast evolution due to the gaining of hypervirulence plasmids allowed this bacterium to cause severe community-transmitted infections in relatively young and healthy hosts since the late 1980s (54, 55). pKP508BN34, with 63.067 bp in size and harboring various mobile elements that contain antimicrobial resistance genes, including *qnrS1* and *bla*_LAP-2_, belongs to the IncFII plasmid group. As seen in the branch highlighted in **Fig. 7B**, most of the strains carrying the closest related plasmids to pKP508BN34 were described in different cities of China – but one in Germany, one in Japan, and three in Thailand –, and were associated with diverse host diseases, such as pneumoniae, pulmonary infection, urinary tract infection, intestinal infection, and diarrhea, according to the data deposited in the bioprojects of NCBI. In some cases, related strains were reported in asymptomatic patients (strains TH164 carrying plasmid pTH164-3 and strain TH114 bearing plasmid pTH114-3, both from Thailand), something reasonable since *K. pneumoniae* is also a member of the gut microbiota (50). To the best of our knowledge, this is the first time that this plasmid is reported in Brazil.

As shown in **Fig. 8A-B**, plasmids pKP98M3N42 and pKP508BN34 are highly structurally conserved between *K. pneumoniae* strains available in the databank. However, plasmid pKP508BN15 (which is present in *K. pneumoniae* 508B) presented a strong structural diversification in the region close to the antibiotic markers, and these changes seem to be related to the activity of the ISSpu21 transposon element located in this region (**Fig. 8C**). Interestingly, most related sequences available in the database are from strains worldwide including some isolated from Asia, Africa, North America, and Europe, but no example of sequences from South America.

**Figure 8.**
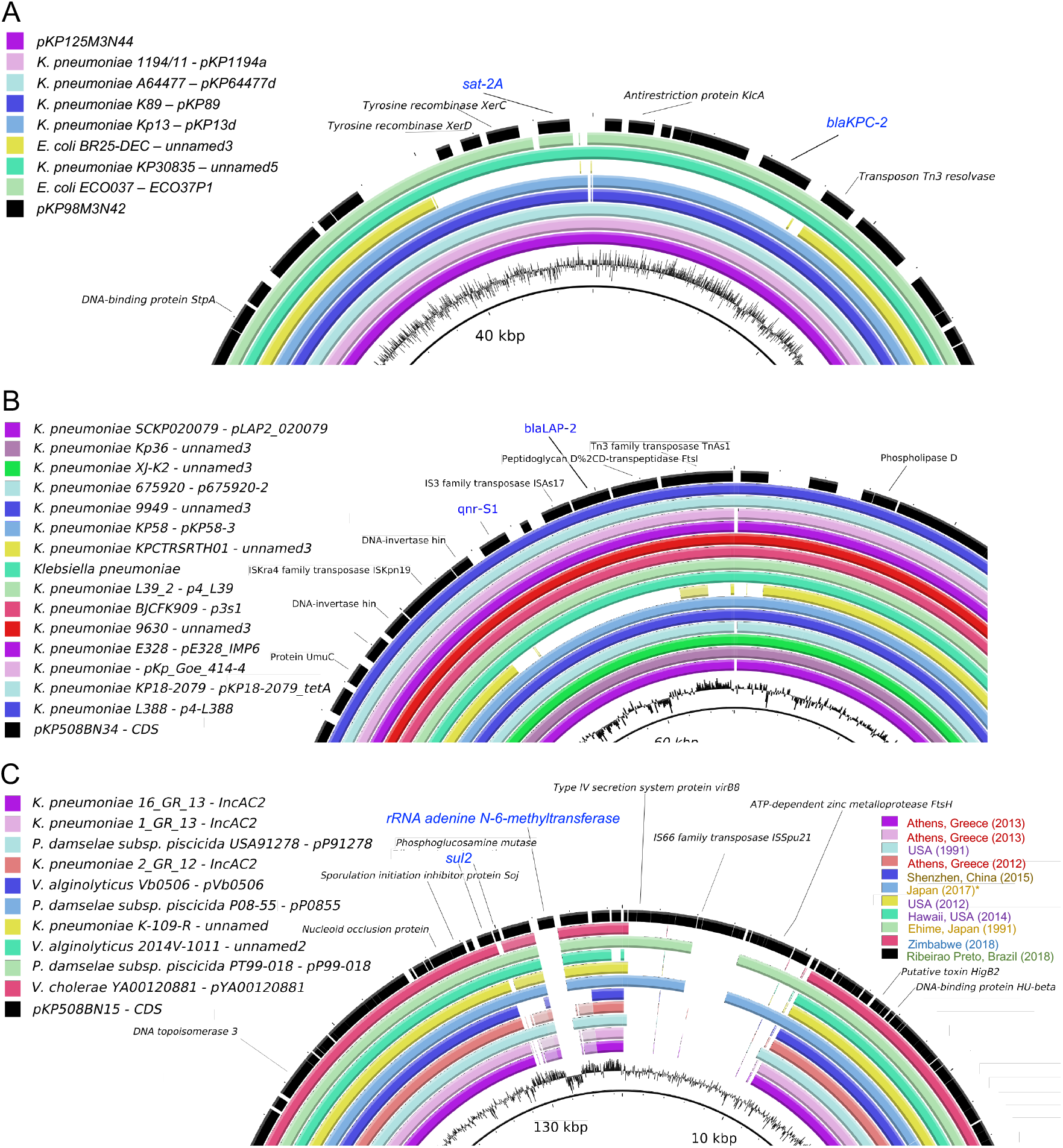
Structural comparison of plasmids from *K. pneumoniae* strains. Nearly identified plasmids were analyzed using blast, and the best hits were used for comparison using BLAST Ring Image Generator (BRIG) (58). For simplification, only divergent regions between the plasmids are shown. **A**) pKP98M3N42 (black) and pKP1253N44 (magenta) are most similar to plasmids found in *K. pneumoniae* and *E. coli,* which have been frequently reported in Brazil (as in Fig. 7A). **B**) pKP508BN34 (black) was most similar to plasmids only isolated from *K. pneumonia,* mostly identified in Asia (shown in Fig. 7B) with a high degree of conservation. **C**) pKP508BN15 (black) was most similar to plasmids found in several different bacterial species (*K. pneumoniae*, *P. damselae subsp. piscicida*, *Vibrio alginolyticus,* and *V. cholerae*) reported in Asia, Africa, and North America. Both *sul2* and rRNA adenine n-6-methyltransferase (*ermC*) resistance markers are located in a highly divergent region of the plasmid and are not present in most related plasmids.

## Conclusions

Here, we sampled clinical bacterial strains to investigate the existence of ARGs and their association with mobile genetic elements. We could identify several ARGs shared between strains from different species, and some of these ARGs are associated with large plasmids, mostly endowed with transposable elements. Instead of focusing on strains from the same species or genus, our approach considered strains co-occurring simultaneously into a hospital setup, aiming to identify shared resistance mechanisms that could have been mobilizing in this environment. While our analysis does not provide unequivocal evidence that these resistance mechanisms are being mobilized among the strains analyzed, we found strongly conserved ARGs located in plasmids and associated with transposon elements that could represent potential mechanisms for the dissemination of antibiotic resistance among clinical strains.

Additionally, by using functional genomics, it was possible to investigate which of the ARGs candidates identified *in silico* could be the main factors associated with the resistance to beta-lactam antibiotics. Furthermore, for many of the bacterial species analyzed here, we could find a low number of available complete genome sequences at the NCBI database. Therefore, while thousand genome sequences are available for classical pathogens (*E. coli*, *K. pneumoniae*, *P. aeruginosa*, etc.), other clinically relevant pathogens such *M. morganii* and *B. cepacia* are underrepresented, which makes challenging to track genomic events associated with the acquisition of pathogenicity elements or resistance mechanisms in hospital-associated infections. Finally, analysis of plasmids from *Klebsiella* strains allowed the identification of both well-known circulating variants in Brazil as new variants that seem to be recently acquired from Asia. Thus, we argue that more systematic efforts should be made to monitor the introduction and propagation of mobile genetic elements harboring ARGs, especially in South America, in order to avoid outbreaks of novel multidrug resistance bacteria.

## Material and methods

### Sample collection

Samples were taken at the Ribeirão Preto Clinics Hospital (HCRP, Ribeirão Preto, Brazil), a tertiary reference hospital in Latin America with 920 beds and 35,000 hospitalizations per year. Samples were isolated from different patients hospitalized at the weeks 44 and 48 of 2018. Strains were obtained from different samples, as indicated in **Fig. 1A**. Thirty-five strains were randomly selected from a total of 105 available and represent different species, which have a major prevalence for *Klebsiella* genus, *E. coli*, *Pseudomonas aeruginosa,* and the genus *Staphylococcus*. After strain characterization by Vitek 2, samples were inactivated and used for genomic DNA extraction and sequencing, as indicated below.

### DNA extraction and genome sequencing

Total genomic DNA was extracted using Wizard Genomic DNA Purification Kit (Promega, Madison, WI, USA) following the manufacturer’s instructions. **Fig. 1B** represents schematically the overall strategy used for WGS analysis. The DNA concentrations were measured fluorometrically (Qubit® 3.0, kit Qubit® dsDNA Broad Range Assay Kit, Life Technologies, Carlsbad, CA, USA). Purified DNA from 35 isolates was prepared for sequencing using Nextera XT DNA Library Prep Kit (Illumina). Libraries were quality assessed using 2100 Bioanalyzer (Agilent Genomics, Santa Clara, CA, USA) and subsequently sequenced using HiSeq 2×150 bp cycle kits (Illumina). On average, 5.5 million reads were generated per library. Adapters were trimmed using Trimmomatic v0.36. Samples were filtered of possible human contamination by aligning the trimmed reads against reference databases using Bowtie2 v2-2.2.3 with the following parameters (-D 20 -R 3 -N 1 -L 20 –very-sensitive-local). Overlapped reads were merged using Flash version 1.2.11. Merged and unmerged reads were assembled using Spades v3.12.0 with the following parameters (-k 21,33,55,77,99,127 --merge). Genome quality (completeness and contamination) was evaluated using CheckM v1.0.7 and QUAST. Genome annotations were performed using Prokka v1.11 with default parameters. Amino acid sequences of all genes identified using Prokka were aligned to the NDARO (National Database of Antibiotic Resistant Organisms) database obtained from NCBI (March, 2020). The alignment was performed using Diamond v0.8.24 with the following parameters (blastx -k 5 -f 6 –E value 0.001). Alignments with >= 60 similarity were selected for further analysis. Quality assessment of sequenced genomes is provided in **Fig. S1**. All genomes are available at NCBI under the BioProject number PRJNA641571.

### Identification of antibiotic resistance genes, plasmids and phylogenomic analysis

The identification of antibiotic resistance genes was performed using the ABRicate pipeline by searching annotated genes using reference databases (ARG-ANNOT, NCBI AMRFinderPlus, CARD and ResFidner), as well as DeepARG (26). The identification of plasmid was performed using Plasmidfinder and contigs harboring ARGs and plasmid related genes were further analyzed in detail. For genomic comparison, reference and assembly genomic data were downloaded from NCBI databank. Phylogenomic analyses were performed using Parsnp and Gingr (51) and phylogenetic trees visualized using iTOL (52).

### Genomic libraries construction

For the cloning of the genomic DNAs in the pSEVA232 (32–34) vector, 2μg of genomic DNA from each strain were digested with Sau3AI. While, pSEVA232 digestion using BamHI and further dephosphorylation were performed. Genomic fragments from 1.5 to 6 kb and the linearized pSEVA232 vector were selected and incubated with the T4 DNA Ligase enzyme in a 2:1 insert/vector ratio. Then, ligations were transformed in the electrocompetent *E. coli* DH10B (53) with a MicroPulser electroporator (Bio-Rad -Hercules, USA). The resulting libraries were analyzed for the percentage of plasmids carrying genomic DNA and the average size of the insert they contained.

### Determination of Minimum Inhibitory Concentrations (MICs)

The MICs were determined in the same culture medium (solid LB) and conditions of the screenings, employing serial dilution of the test antibiotics (Amoxicillin, Oxacillin, or Penicillin G). Solid medium plates supplemented with kanamycin (50 μg mL^−1^), IPTG (100 μM) and dilutions of each beta-lactam antibiotic were inoculated with approximately 2.5 × 10^6^ CFUs of *E. coli* DH10B harboring the pSEVA232 vector. We performed dilutions of antibiotics and culture media according to the CLSI M100 supplement (54).

### Screening and phenotype confirmation

Three pools of clones (2.5 × 10^6^ clones per plate) were plated from each library on solid LB supplemented with kanamycin (50 μg μL^−1^), IPTG (100 μM) and inhibitory concentrations of each beta-lactam antibiotic (8 μg mL^−1^ for amoxicillin, 32 μg mL^−1^ for penicillin G or 256 μg mL^−1^ for oxacillin). Positive clones were cultured in liquid LB medium supplemented with kanamycin (50 μg mL^−1^) for extraction of plasmid DNA with the Wizard Plus SV Minipreps DNA Purification System Kit (Promega - Madison, USA). Plasmids with unique EcoRI/HindIII restriction patterns were subjected to re-transformation and phenotypic confirmation by streaking the clones in solid LB medium supplemented with kanamycin (50 μg mL^−1^), IPTG (100 μM) and the test antibiotics.

### Extraction of insert sequence from assembled genomes

Plasmid DNA was sequenced on both strands by primer walking using the ABI PRISMDye Terminator Cycle Sequencing Ready Reaction kit (PerkinElmer) and an ABI PRISM 377 sequencer (Perkin-Elmer) according to the manufacturer’s instructions. A python script was developed to extract and annotate inserts from Sanger reads, which is available on GitHub. First, the algorithm converts .ab1 files to the fasta format and searches for the read sequence in the assembled genome using BLAST+ (55). We then extract the inter-read region to a fasta file, which is then annotated by PROKKA (56).

## Acknowledgments

The authors are thanks to lab colleagues for their insightful comments and suggestions throughout the course of this study. We thank to Murilo Henrique Anzoline Cassiano for his initial support on genome assembly. We also thank to Dr. Roberto Martinez for providing the bacterial strains used in this study.

---

**Figure S1.**
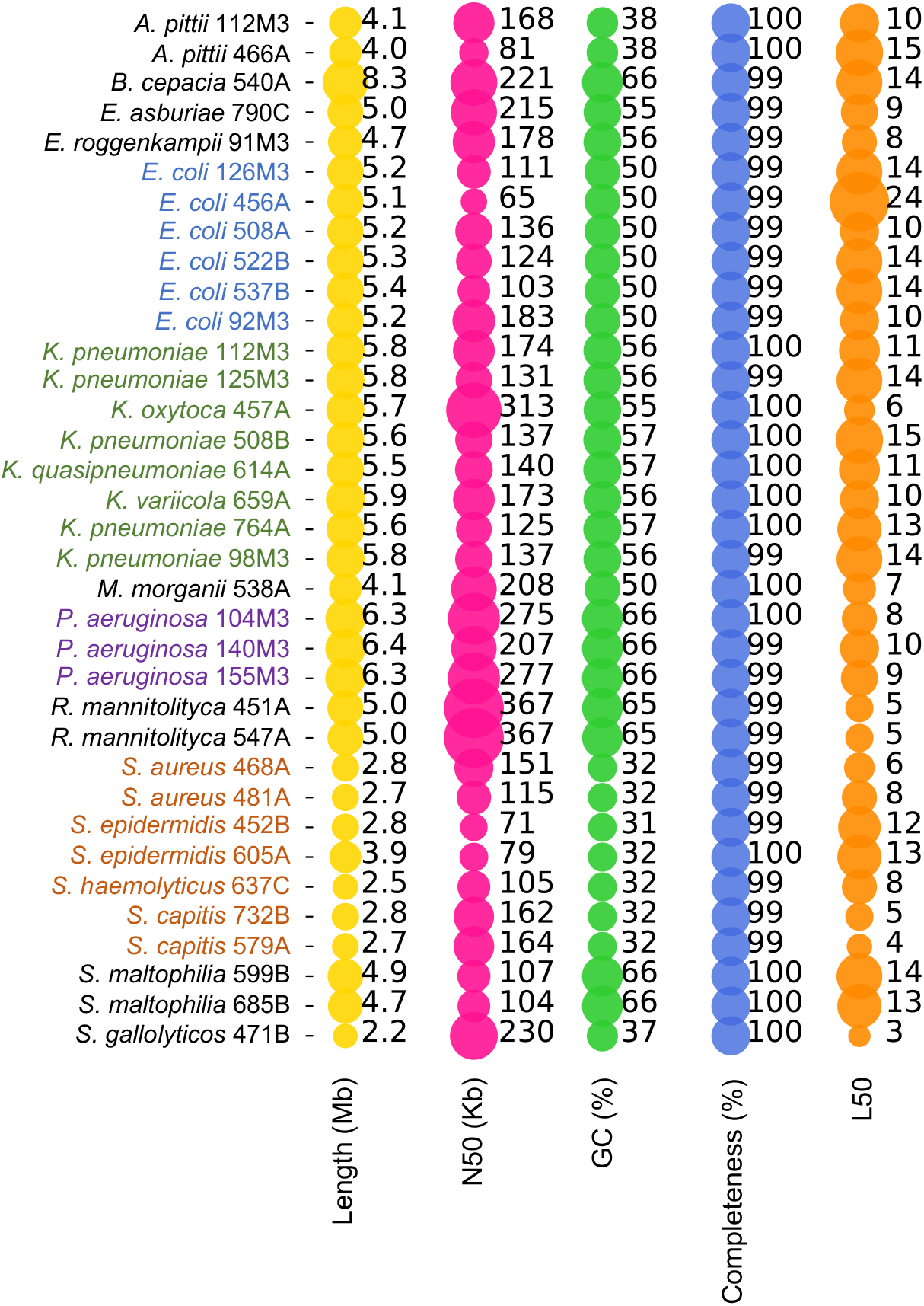
General features of genomes sequenced in this study.

**Figure S2.**
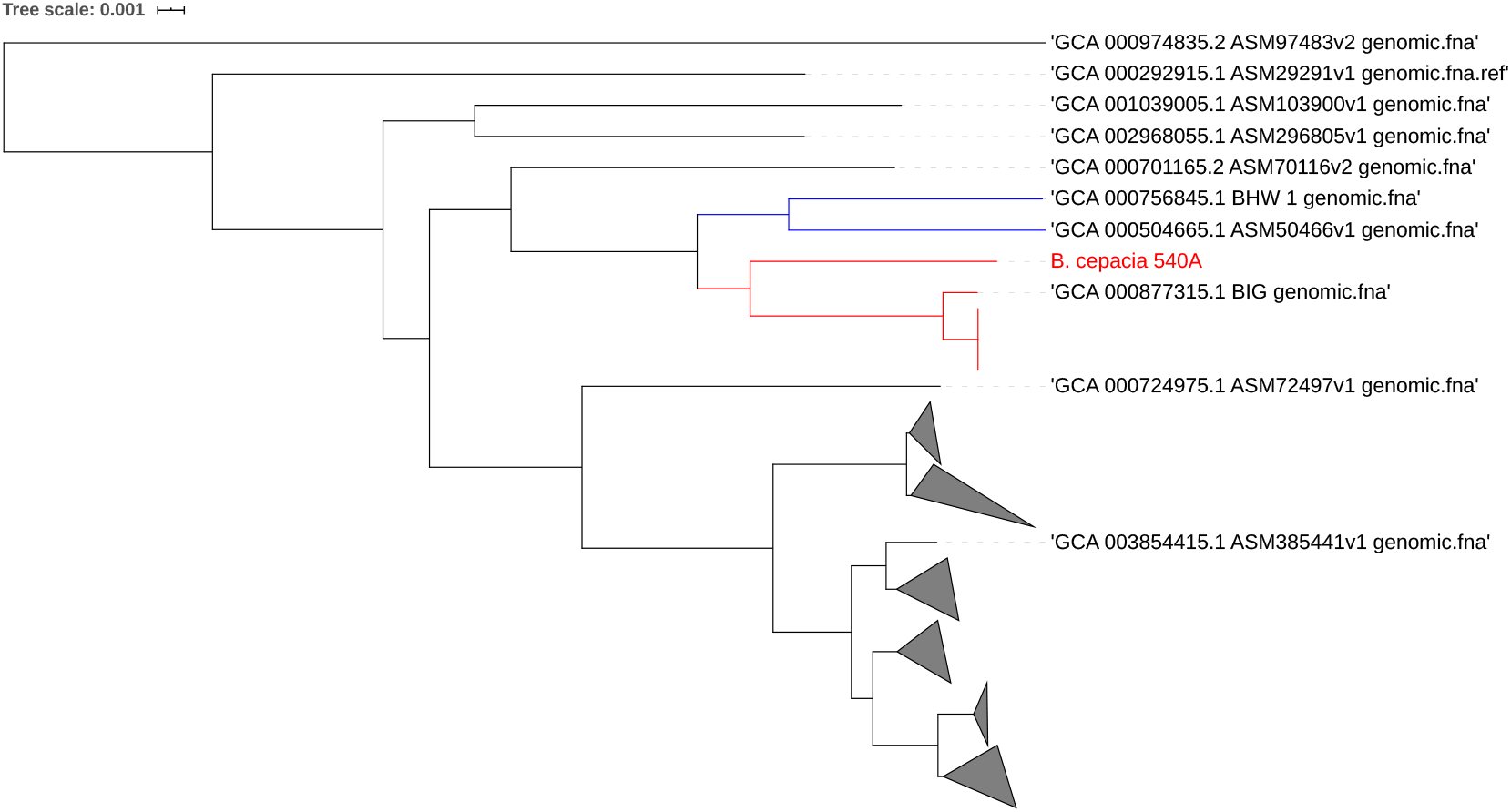
Phylogenetic tree of *B. cepacia* 540A and total number of available *B. cepacia* genomes (166) in NCBI at March 2020. For simplicity, branch with high similarity were compressed (gray triangles). NCBI accession number are indicated for *B. cepacia* strains compared in the figure. Blue branches are two *B. cepacian* endophytic strains isolated in Australia, while red branches are *B. cepacian* isolated from patients.

**Figure S3.**
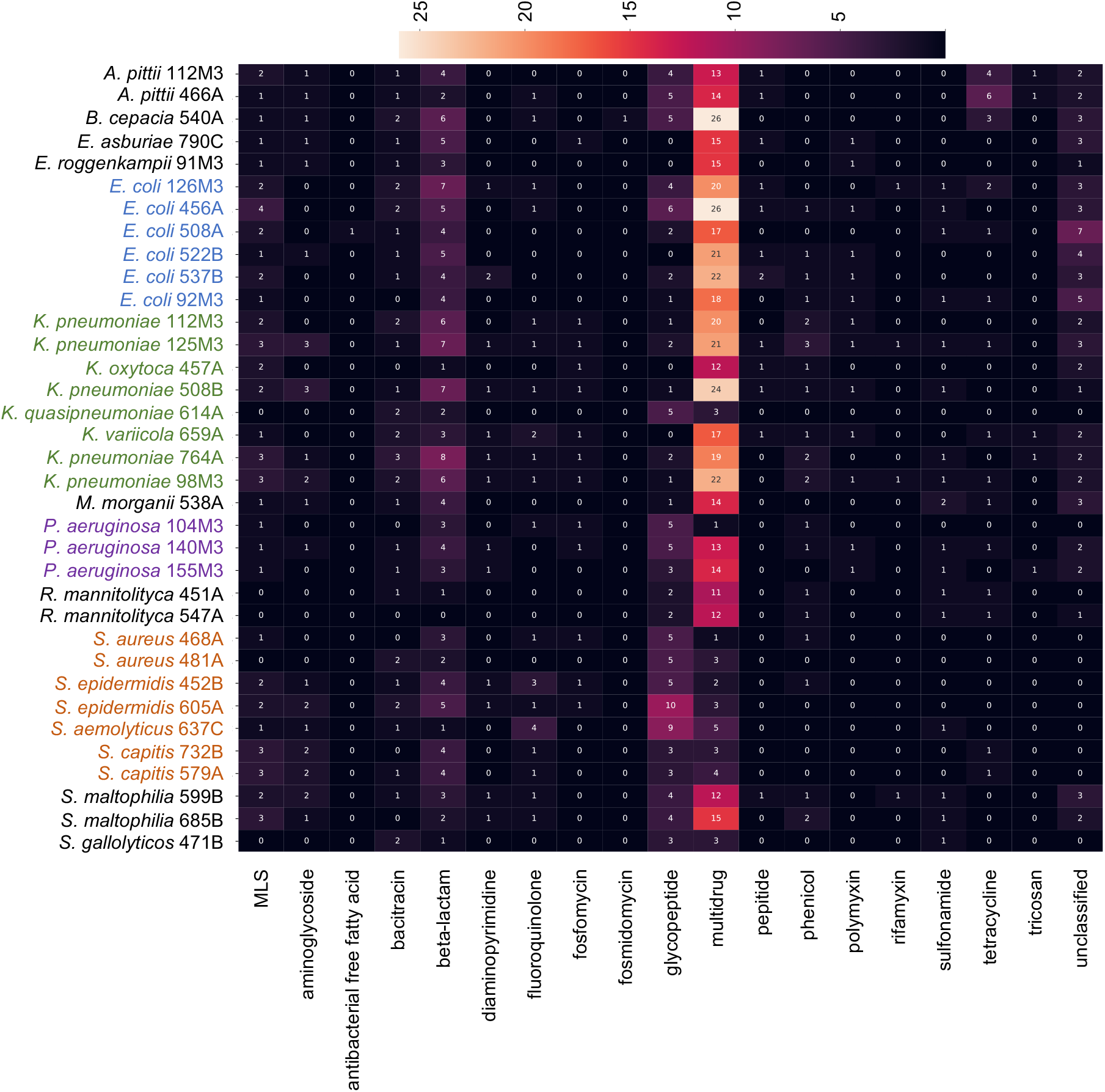
Heatmap showing the presence of ARGs identified by DeepARG in nearly sequenced genomes.

**Figure S4.**
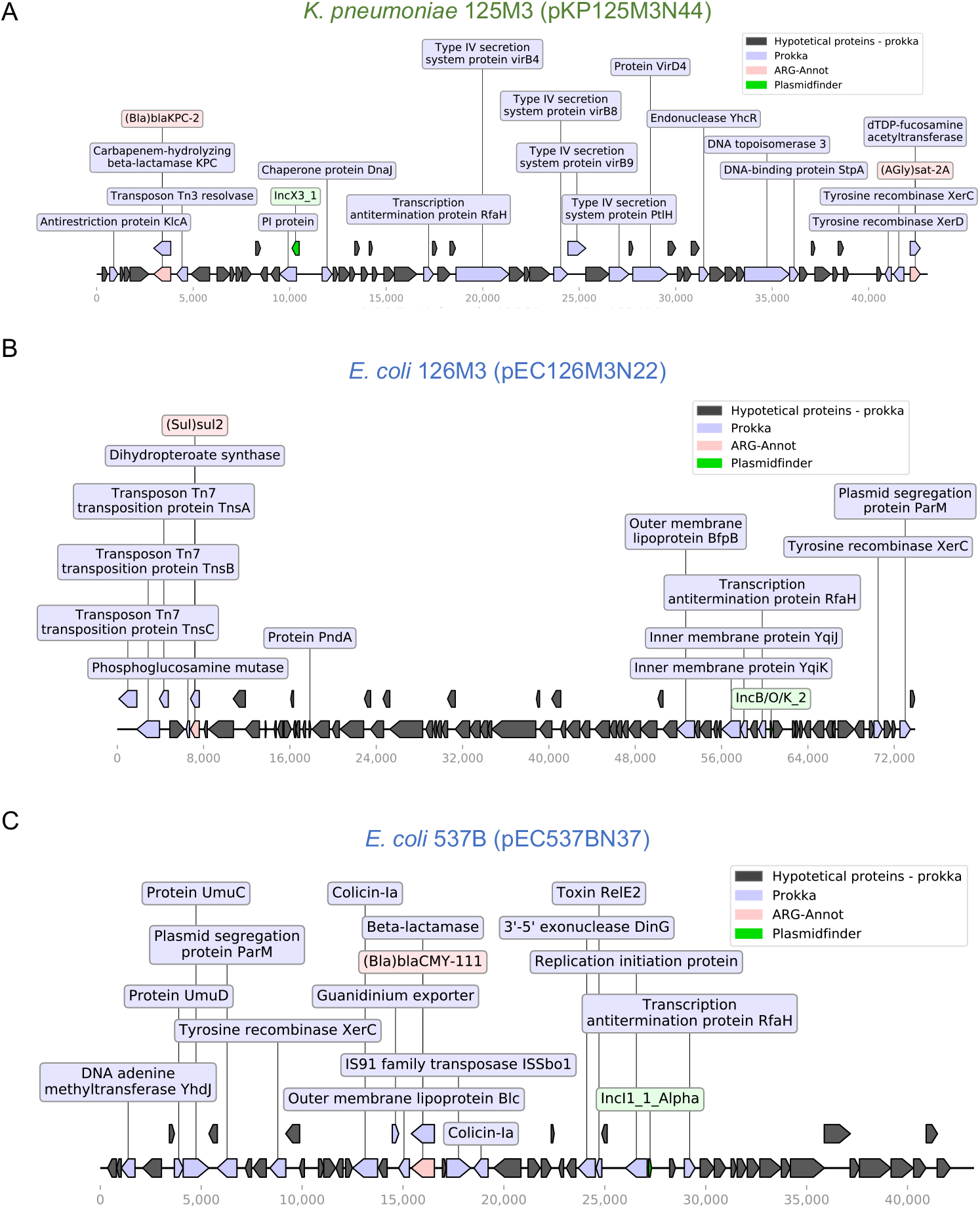
Schematic representation of genes in identified plasmids exposing the coexistence of ARGs with transposon elements. **A**) Plasmid pKP125M3N44 (43 kb) from *K. pneumoniae* 125M3. This plasmid is identical to plasmid pKP98M3N42 from *K. pneumoniae* 98M3 (Fig. 4A). **B**) Plasmid pEC126M3N22 (73,9 kb) from *E. coli 126M3*. **C**) Plasmid pEC537BN37 (43,3 kb) from *E. coli 537B.* Legends represents the colors code for the identified genes.

**Figure S5.**
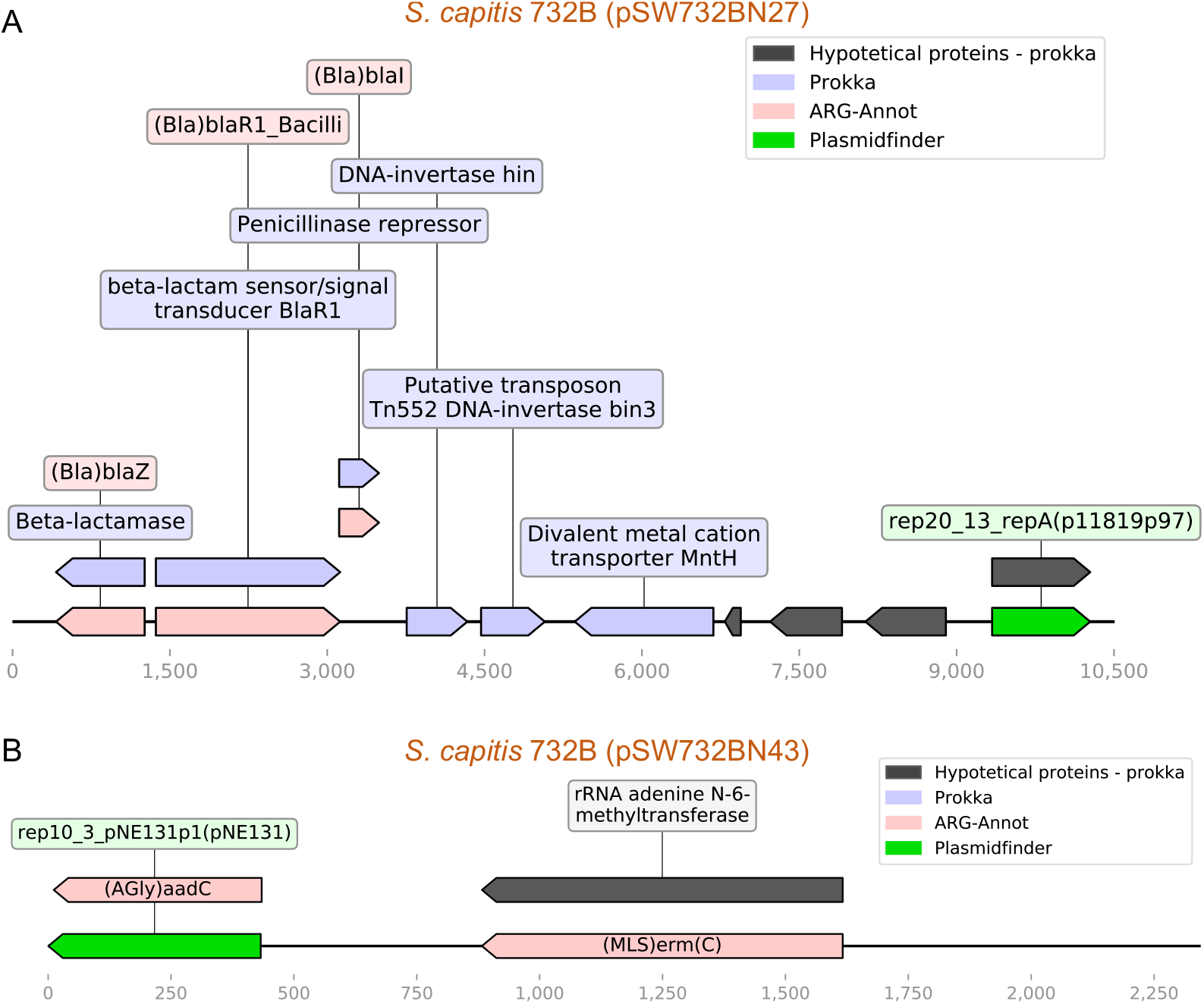
Schematic representation of plasmids identified in *Staphylococcus* strains showing the coexistence of ARGs with transposon elements. **A**) Plasmid pSW732BN27 (10.5 kb) from *S. capitis 732B*. **B**) Plasmid pSW732BN43 (~2.3kb) from *S. capitis 732B*.

